# Explainable Multimodal Prognostic Modeling Uncovers New Clinically Relevant Pathological Features in a Multi-Institutional Pancreatic Neuroendocrine Tumor Cohort

**DOI:** 10.64898/2026.11.02.763764

**Authors:** Arrun Sivasubramanian, Mohammad Salimian, Jithin Reddy Vennapusa, Lucie Dequiedt, Yu Shen, Valentina Matos-Romero, Yutong Zhu, Uwakmfon-Abasi Ebong, Zoe Gaillard, Ian Reucroft, Hyun-Nam Yun, Seung-Mo Hong, Jinho Shin, Atsuko Kasajima, Günter Klöppel, Jin He, Ralph H. Hruban, Ashley L Kiemen

**Affiliations:** The Sol Goldman Pancreatic Cancer Research Center, Department of Pathology, Johns Hopkins University School of Medicine, Baltimore MD, USA; Bloomberg School of Public Health, Johns Hopkins University, Baltimore, MD, USA; Chemical & Biomolecular Engineering, Johns Hopkins University, Baltimore, MD, USA; Department of Pathology, Asan Medical Center, University of Ulsan College of Medicine, Seoul, Republic of Korea; Institute of Pathology, Technical University of Munich School of Medicine and Health, München, Germany; Department of Surgery, Johns Hopkins University, Baltimore MD, USA; Department of Oncology, Johns Hopkins University, Baltimore MD, USA; Department of Functional Anatomy & Evolution, Johns Hopkins University, Baltimore MD, USA

**Author notes:** To whom correspondence should be addressed. Ashley Kiemen, Ph.D. 1729 East Monument Street Room 10219 The Johns Hopkins University School of Medicine Baltimore, MD 21287 Ralph H. Hruban, M.D. Weinberg Room 2267 401 North Broadway The Johns Hopkins University School of Medicine Baltimore, MD 21231.

**Keywords:** Pancreatic neuroendocrine tumor, Well-differentiated neuroendocrine tumor, Digital pathology, Risk prediction, Tumor stroma

## Abstract

Although many patients with a well-differentiated pancreatic neuroendocrine tumor (PanNET) are cured after surgical resection, it is hard to predict outcome in these patients. Current prognostic assessment relies largely on Ki-67 proliferation index, tumor stage, and established adverse pathologic features, such as vascular and perineural invasion. However, individually, these variables incompletely explain why patients with similar clinicopathologic profiles sometimes experience markedly different outcomes. To address this challenge, we assembled three independent cohorts with a surgically resected PanNET. In total, we included 1,438 patients with approximately 30,000 histology images, and approximately 12,000 person-years of follow-up. We separately modeled structured clinicopathologic information with Cox regression and tumor-centered pathology images with deep learning on an internal cohort treated at Johns Hopkins Hospital and performed multimodal continuous-time survival modeling to reveal which clinical and pathologic features are most important in predicting patient outcome.

Across Cox models of censored time to recurrence-free survival, a text-based clinicopathological model that incorporated known prognosticators achieved a C-index of approximately 0.87, the image-based histology model approximately 0.80, and decision-level late multimodal fusion approximately 0.88 on the internal test set. The clinical model demonstrated strong prognostic discrimination using established risk factors, mainly relying on the TNM stage and Ki-67 index. Primary tumor histology retained substantial prognostic signal, with analysis restricted to tumor-centered regions to reduce reliance on unrelated tissues that might indirectly indicate disease extent. Interpretation of high-attention image tiles revealed regions of importance in the primary tumor, including recognized adverse morphologic features such as the leading edge of the tumor and foci of vascular or perineural invasion. Unexpectedly, intratumoral and peritumoral stroma accounted for nearly 50% of the model’s high-attention regions, suggesting that the stroma may yield equal prognostic information to neoplastic cells in recurrence-prediction models.

To test generalizability, we further evaluated the trained models in independent cohorts from two institutions with different demographics, where discrimination varied by site (mean late-fusion C-indices of 0.864 and 0.723). Together with the improved discrimination achieved through late fusion, our findings suggest that current clinicopathologic variables can be modeled to form reliable prognosticators, and that routine hematoxylin and eosin (H&E) stained slides of the primary tumor contain additional prognostically important information not fully captured by conventional clinicopathologic assessment, such that together, the two sources provide complementary information.

## INTRODUCTION

Well-differentiated pancreatic neuroendocrine tumors (PanNETs) are malignant neoplasms of the pancreas. Many patients with a PanNET are cured by surgical resection, however, a clinically important subset recur years after surgery.^1–5^ Established prognosticators for patients with a surgically resected PanNET include TNM stage, Ki-67 proliferation index, and vascular and perineural invasion.^6–10^ Routine histopathology of primary tumors may contain additional prognostic information that is not fully captured by these predefined variables, including the morphology of the invasive front (circumscribed versus invasive), architectural heterogeneity, and rare aggressive foci.^11–14^ This raises the possibility that other prognostic information in primary tumors, not appreciated by pathologists, may also contribute to patient outcome.

Computational pathology has created a practical framework to address this challenge.^15,16^ Pathology foundation models encode morphology from hematoxylin and eosin (H&E) images into transferable numerical representations,^17–24^ while language encoders can similarly represent standardized clinicopathologic statements.^25–28^ Multimodal methods have shown that H&E representations can be integrated with clinical or molecular information for survival modeling using vision-language architectures and multiple instance learning (MIL) ^29–39^ These patient-level representations can then be used in Cox proportional hazards models to estimate relative risk from observed event times while accounting for censored follow-up.^40^ Together, these approaches create an opportunity to compare the prognostic information contained in routinely recorded clinicopathologic variables with the richer spatial information preserved in whole-slide histology, and to determine whether the two sources provide complementary information.

PanNET presents a particularly challenging setting for prognostic modeling, as the rarity of the disease limits the number of cases available for analysis, and prolonged patient survival requires long patient follow-up.^41–46^ Practical challenges include patient histories span multiple decades across institutions: clinical documentation that varies over these timeframes, and pathology slides are prepared with heterogeneous fixation, staining, and scanning protocols.^47–49^ Here, we assembled three independent cohorts, from universities, with linked clinicopathologic records, high-resolution H&E whole-slide images (WSIs), and postoperative follow-up. We integrated clinical records, linked them to digitized WSIs, and defined time-to-event outcomes that retained both the timing of recurrence or disease-related death and the duration of available follow-up. In total, 1,438 patients diagnosed with PanNET we included in the retrospective study. We developed models on patients treated at the Johns Hopkins Hospital (JHH) and internally benchmarked them, and then externally validated the models on independent cohorts from the Technical University of Munich Hospital (TUM), Germany, and Asan Medical Center (AMC), Korea. We trained two models. The first predicted disease-free recurrence in all eligible patients for whom pathology and long-term follow-up were known. The second focused on the most clinically challenging patients: those who presented with localized, non-metastatic, disease and who underwent pancreatic resection, excluding those with distant metastasis whose negative prognosis was clear and excluding those who underwent enucleation, whose positive prognosis was clear.

We assessed the individual and combinatorial prognostic value of established clinicopathologic variables with Cox models and examined whether their associations with outcome remained informative across institutions. We then modeled primary-tumor H&E morphology using foundation-model tile embeddings and attention-based multiple instance learning (ABMIL) to assess whether image-derived features provided additional prognostic information. By mapping tile attention scores back to their original locations in each patient’s primary tumor and incorporating pathologist review, we were able to determine the histologic features the model emphasized, such as the tumor center, leading edge, and stromal compartments. Finally, we examined three ways to integrate clinicopathologic and histologic information: fixed-weight fusion of unimodal risk scores, stacked cox modeling of those scores, and learned feature-level fusion. Using matched patient-level test folds, we aimed to determine whether these strategies improved postoperative risk stratification for patients with a surgically resected PanNET and to assess their generalizability in independent cohorts.

## RESULTS

### Construction of longitudinal multimodal PanNET cohorts for outcome-defined risk modeling

The retrospective study utilizes three cohorts: one internal used for training and validation, and two for external validation. The internal cohort consisted of a clinically annotated cohort of well-differentiated PanNETs surgically resected from 903 patients treated at JHH between 1980 and 2025. We digitized a total of 26,625 archival whole slides and curated 3,888 H&E images containing the primary tumor. For each case, we harmonized clinicopathologic records and long-term postoperative follow-up^50^ and digitized WSIs. Redundant patient visits or duplicate records, cases with insufficient (<5 years) or unresolved follow-up, deaths not related to PanNET progression, and patients treated with neoadjuvant therapy were excluded to ensure the integrity of our prognostic model. Two independent PanNET cohorts, one from the TUM with 207 patients, and the second from AMC with 328 patients, were assembled for external validation using the same selection criteria as JHH (Fig. 1a)

**Figure 1.**
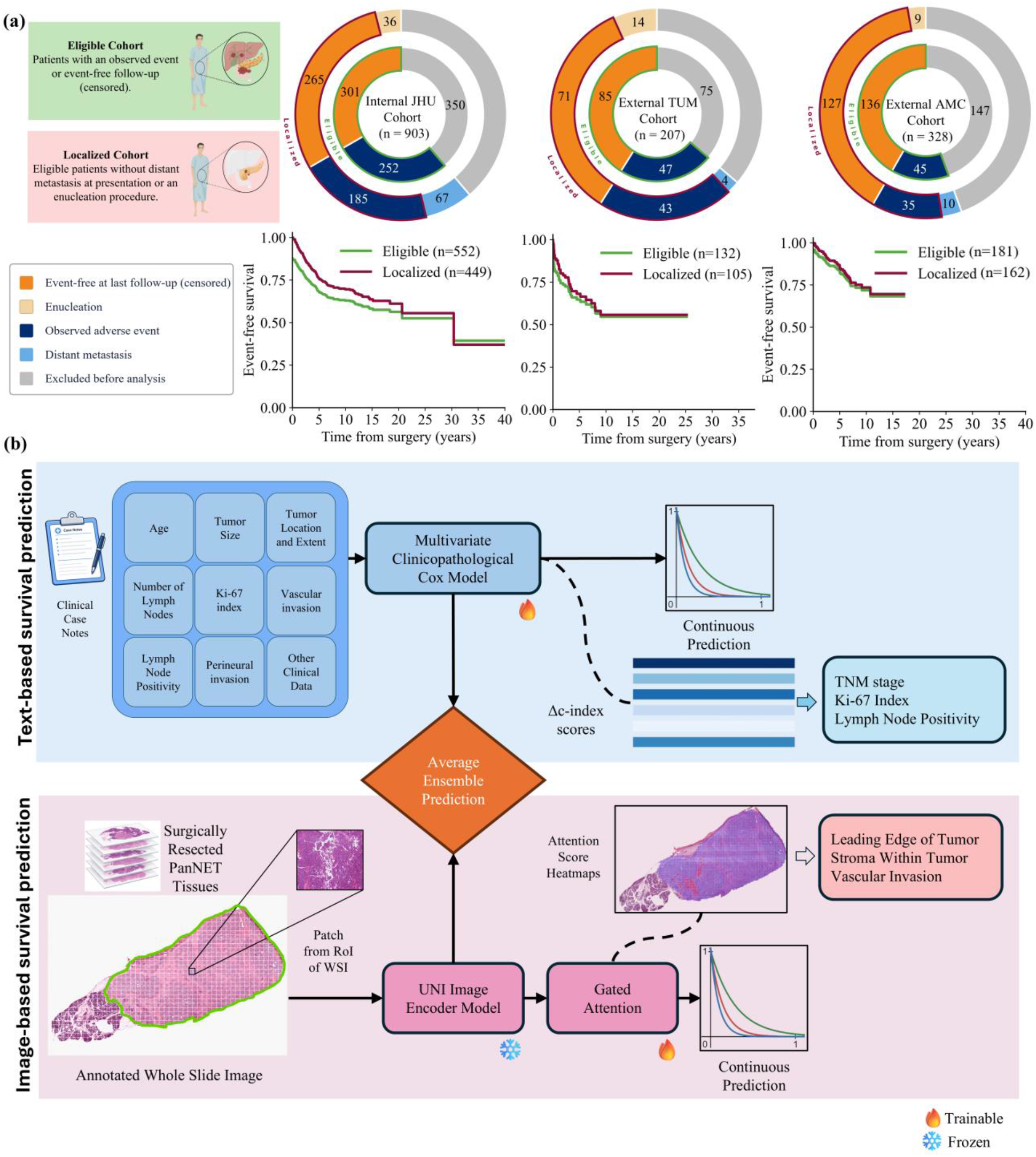
Study cohorts, and overall framework for multimodal survival prediction. **a)** Concentric diagrams show the eligible (inner ring) and localized (outer ring) cohorts at Johns Hopkins Hospital (JHH), the Technical University of Munich (TUM) and Asan Medical Center (AMC), alongside the Kaplan-Meier estimates at each institution. **b)** Overview of the modeling framework.

For the three independent PanNET datasets, we defined the eligible cohort: consisting of all patients for whom we possessed digitized histology of their primary tumor and had information regarding either a documented recurrence or at least five years of recurrence-free follow-up. For the eligible patients, the patient’s minimum time to recurrence or death was chosen as the censor for modeling the continuous time-to-event Cox proportion hazard model. To further test our workflow in a more clinically meaningful population, we defined the localized cohort: consisting of patients who did not have distant metastases at the time of surgery. We also excluded patients with extremely indolent diseases who only underwent enucleation. The slopes of the Kaplan-Meier curves showed that metastatic presentation was associated with markedly worse outcomes, whereas enucleation was associated with extremely favorable outcomes,^50^ motivating their exclusion from the set of patients with localized disease. This sub-analysis therefore focused on patients for whom postoperative risk stratification is most valuable.

For each case, clinical and pathologic variables were converted into standardized fields, and all tumor-bearing H&E-stained WSIs of the primary tumor were selected for image analysis (Fig. 1b). The resulting cohort provided a rare longitudinal multimodal resource for PanNET prognosis via evaluation of both clinicopathological variables from case notes and primary tumor morphology.

### Clinical case-note modeling identifies outcome-associated clinical variables

For each patient, we harmonized clinical case notes and pathology reports into 14 structured clinicopathologic variables, including TNM stage, Ki-67 index, lymph-node involvement, and vascular and perineural invasion.^51–53^ Continuous variables were represented using cubic B-splines to allow nonlinear associations with outcome, whereas categorical variables were converted into one-hot encoded vectors (Fig 2a). These fields were used to model observed time to event while retaining patients with censored follow-up on the internal training cohort and then evaluated on a held-out test set and the external validation cohorts.

**Figure 2.**
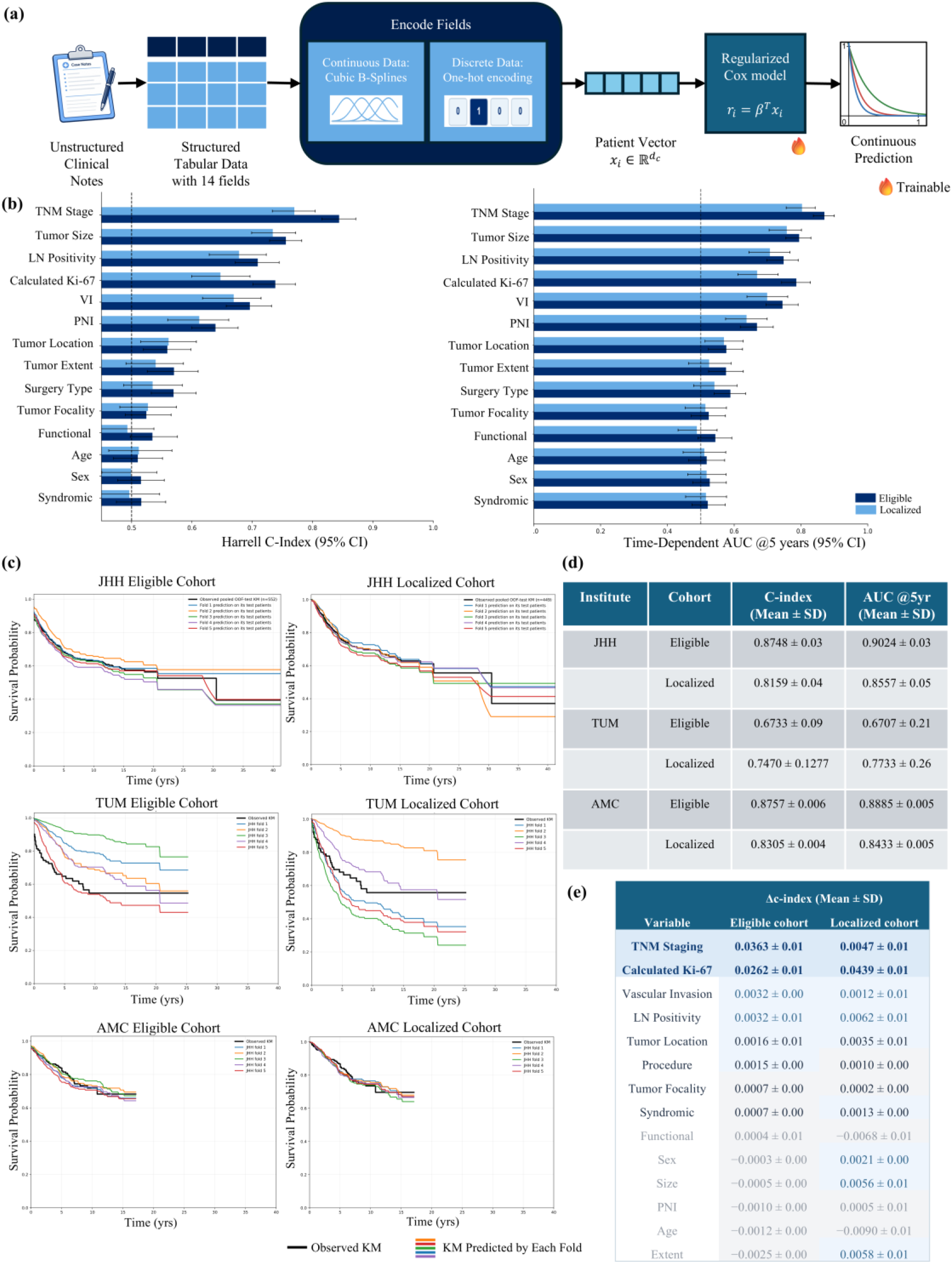
Structured clinical features predict outcomes for patients with a PanNET across internal and external cohorts. **a)** Clinical case notes and pathology reports were harmonized into 14 patient-level variables and passed to the clinicopathological model. **b)** Held-out univariate discrimination for each variable in the JHH cohorts, measured by Harrell’s C-index and five-year time-dependent AUC (error bars indicate 95% confidence intervals). **c)** Observed Kaplan-Meier survival (black) and mean predicted survival from each of five Cox models (colored) in JHH, TUM and AMC. **d)** C-index and five-year time-dependent AUC, reported as mean ± SD across the five models or held-out folds. **e)** Variable contribution measured as the change in held-out C-index after removing each variable and refitting the multivariable model.

Using the JHH cohort, we first fitted a univariate Cox model for each variable and evaluated predictions in held-out patient folds. In the eligible cohort of 552 JHH patients, TNM stage had the highest pooled out-of-fold C-index (0.844; 95% CI, 0.816-0.872) and a five-year time-dependent AUC of 0.870. Tumor size (C-index, 0.756; 95% CI, 0.729-0.782), Ki-67 index (0.738; 0.701-0.772) and the positive-to-harvested lymph-node ratio (0.709; 0.672-0.745) also discriminated outcomes. In the cohort of 449 JHH patients with localized disease, TNM stage remained the strongest individual variable (0.770; 0.733-0.804), followed by tumor size (0.734; 0.699-0.772). These findings are consistent with the established prognostic relevance of tumor burden and proliferative activity in resected PanNETs.^54,55,54,55^

We next combined all 14 variables in a regularized multivariable Cox model. In the JHH cohort, across the five held-out test folds, the mean C-index was 0.8748 ± 0.034 in the eligible JHH cohort and 0.8159 ± 0.042 in the localized JHH cohort; mean five-year AUCs were 0.9024 ± 0.034 and 0.8557 ± 0.053, respectively. Each patient appeared in one test fold. The observed survival curve and the five curves summarizing predictions for the respective held-out folds are shown in Fig. 2c. The model thus retained strong prognostic discrimination using the grading and staging variables of routine clinical practice in both eligible and localized cohorts.

We applied each of the five trained JHH models, without retraining, to the independent TUM and AMC cohorts (Fig. 2c). In TUM, the eligible cohort had a mean C-index of 0.6733 ± 0.09 and five-year AUC of 0.6707 ± 0.21; the localized TUM cohort reached 0.7470 ± 0.1277 and 0.7733 ± 0.26, respectively. For the AMC cohort, the corresponding C-indices were 0.8757 ± 0.006 and 0.8305 ± 0.004, with five-year AUCs of 0.8885 ± 0.005 and 0.8433 ± 0.005. These values are means and standard deviations across the five JHH models, each evaluated on the same external patients. The results demonstrate strong transfer to AMC and prognostic discrimination in TUM, with greater variation between models in the latter cohort (Fig. 2d).

Finally, we assessed each variable’s contribution in the internal validation cohort by removing a variable, refitting the combined model and measuring the change in held-out C-index (Fig. 2e). TNM stage and Ki-67 index produced the largest mean reductions in the eligible cohort (0.0363 and 0.0262, respectively). Ki-67 index produced the largest reduction in the cohort with localized disease (0.0439). Together, the univariate and multivariable analyses reinforce the clinical value of recording stage and proliferative activity and establish a strong clinicopathologic reference for evaluating image-derived prognostic information.

### Histology-based prognostic modeling of the primary tumors highlights tumor-associated stroma as a novel prognostic indicator in PanNET

We next evaluated whether the morphology of the primary tumor could predict postoperative disease status. Rather than evaluate the entire digitized histology corpus, which contained a variable mix of slides from the primary tumor, metastatic sites, and non-neoplastic tissues such as spleen duodenum, lymph nodes, depending on the surgery type and tumor stage, we restricted our model to the immediate vicinity of the primary tumor cells in WSIs To do this, we developed a two-step tumor localization strategy to exclude non-target regions. ^56^We first applied CODAVision,^56^ a model for semantic segmentation of tumor cells, to segment twelve pancreatic tissue compartments including PanNET in each WSI, and then spatially constrained the PanNET segmentation with pathologist-guided coarse annotation of the primary tumor to remove from the analysis tumor tissue in metastatic sites such as lymph nodes and duodenum. (Supplementary Fig. S1).

The intersection between the clinician-guided outline and the CODAVision PanNET mask defined the final primary tumor region of interest (RoI). This careful subsampling eliminated the possibility that the model spuriously learned from the tissue context. For example, only distal pancreatectomy surgeries contain splenic tissue and hold the possibility of tumor invasion into the spleen. Therefore, the presence of splenic tissue in a microscope slide would inform the computer vision model of the surgery type and possibly local aggressive behavior. ^57–60^ We dilated the resulting tumor ROI by five tile widths (approximately 515 µm) to retain the tumor together with its surrounding microenvironment. We extracted UNI2 ^18^ tile embeddings from this ROI on all cohorts, aggregated using ABMIL and trained with a Cox survival objective using just the JHH cohort (Fig. 3a), and evaluated the trained model on the external cohorts.

**Figure 3.**
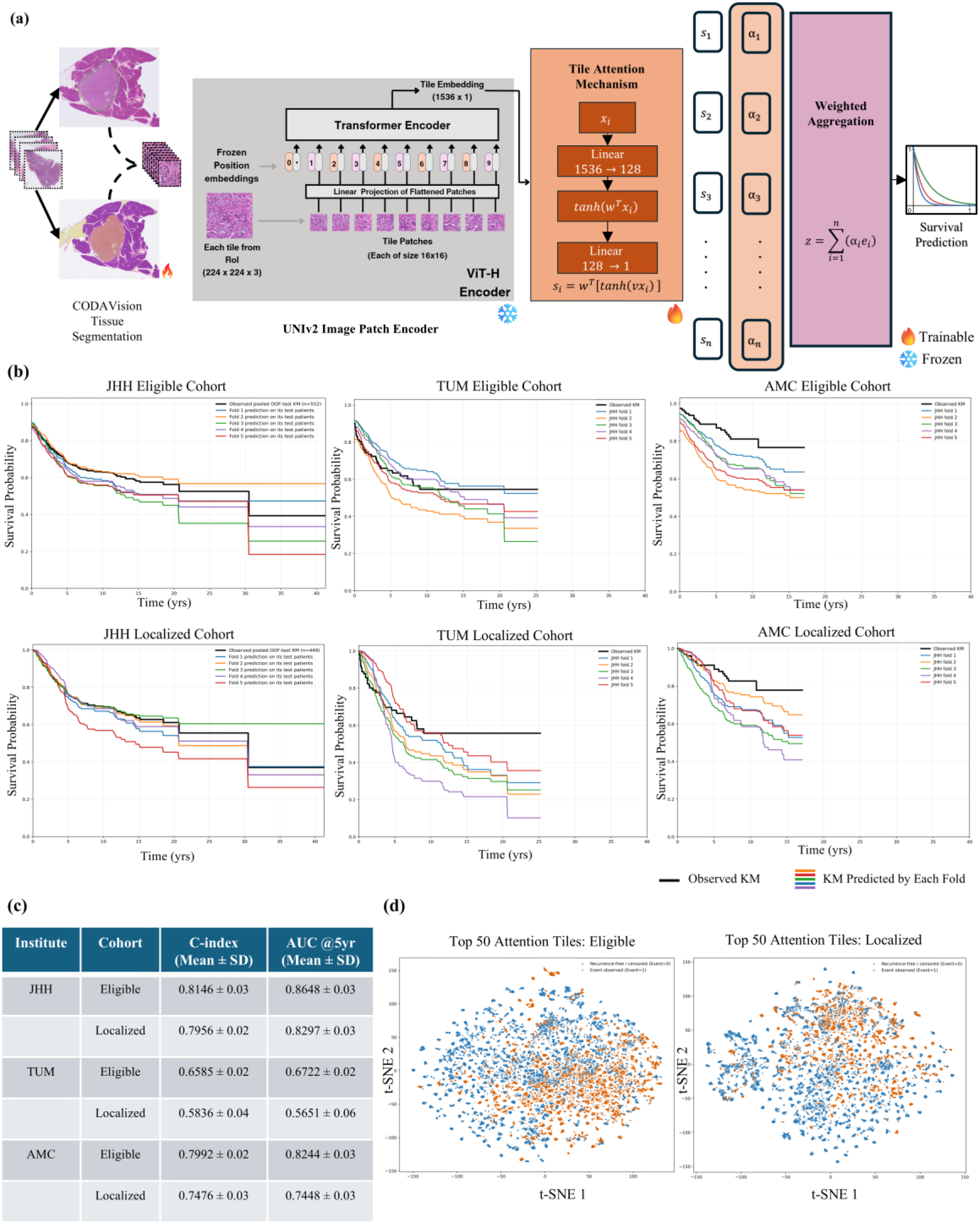
Tumor-region histology predicts postoperative outcomes. **a)** Pathologist-guided primary-tumor localization combined with CODAVision segmentation defines the region of interest. Tumor-region tiles are encoded by frozen UNI2 and pooled by attention-based multiple instance learning to estimate patient-level Cox risk. **b)** Observed Kaplan-Meier survival (black) and survival predicted by each of five-fold-specific models (colored) in eligible and localized cohorts at JHH, TUM and AMC. **c)** Harrell’s C-index and five-year time-dependent AUC, shown as mean ± SD. across held-out folds at JHH and across five JHH models applied to the same patients at each external institution. **d)** Two-dimensional t-SNE projections of the highest-attention tile embeddings, colored by whether an event was observed during follow-up.

Observed Kaplan-Meier survival and mean survival predicted for the patients in each held-out fold are shown in Fig. 3b. In JHH held-out testing, the image model achieved a mean C-index of 0.8146 ± 0.03 and a mean five-year time-dependent AUC of 0.8648 ± 0.03 in the eligible set of patients. Corresponding values in the localized set of patients were 0.7956 ± 0.02 and 0.8297 ± 0.03 (Fig. 3c). These results demonstrate substantial prognostic information in primary-tumor histology without clinical variables.

External tile embeddings showed institution-associated differences from the JHH training distribution. Feature-wise mean and standard-deviation alignment brought the external embeddings closer to the JHH distribution in the visualized t-stochastic nearest embedding (T-SNE) projections (Supplementary Fig. S2). Following physical-scale harmonization and embedding alignment, we applied the five JHH image models to the TUM and AMC cohorts without fine-tuning. In the TUM cohort, mean C-indices were 0.6585 ± 0.02 for the eligible cohort and 0.5836 ± 0.04 for the localized cohort, with corresponding five-year AUCs of 0.6722 ± 0.02 and 0.5651 ± 0.06. In the AMC cohort, mean C-indices were 0.7992 ± 0.02 and 0.7476 ± 0.03, with AUCs of 0.8244 ± 0.03 and 0.7448 ± 0.03. Thus, image-derived prognostic discrimination transferred more strongly to AMC than to TUM, particularly for localized disease.

Finally, we projected the embeddings of each patient’s 50 highest-attention tiles into two dimensions using T-SNE. (Fig. 3d). Tiles from patients with observed events and those with event-free censored follow-up showed distinct grouping tendencies, particularly in the localized cohort, despite some overlap. This pattern supports the presence of outcome-associated morphologic differences within the regions emphasized by the model.

To identify the regions of the primary tumor responsible for the model’s predictions, we generated attention-score heatmaps overlaid on the original WSIs. To move beyond qualitative inspection, we developed a pathologist-facing graphical user interface in which a board-certified anatomic pathologist evaluated the relative importance of predefined tumor-relevant regions for each WSI. With this interface we were able to classify areas identified in the heat maps into one of seven predefined regions: tumor center (non-leading edge of the tumor), the leading edge of the tumor, intratumoral stroma, peritumoral stroma, stroma away from tumor, vascular invasion, and perineural invasion. We additionally allowed free-text entry for instances where model attention focused somewhere other than the seven predefined areas. This review showed that high-attention regions frequently localized to biologically plausible compartments, particularly, leading-edge regions (Fig 4a.), tumor-stroma interfaces (Fig 4b.), intratumoral or peritumoral stroma, and foci suggestive of vascular or perineural invasion. Combined selections of attention to the tumor center and leading edge were similar, and account for 47.3% and 43.2% of the regions identified, respectively (Fig. 4c). Together with the top-tile embedding analysis, this review suggests that the image model drew on both the tumor morphology and the tumor’s stromal microenvironment when predicting disease-free survival.

**Figure 4.**
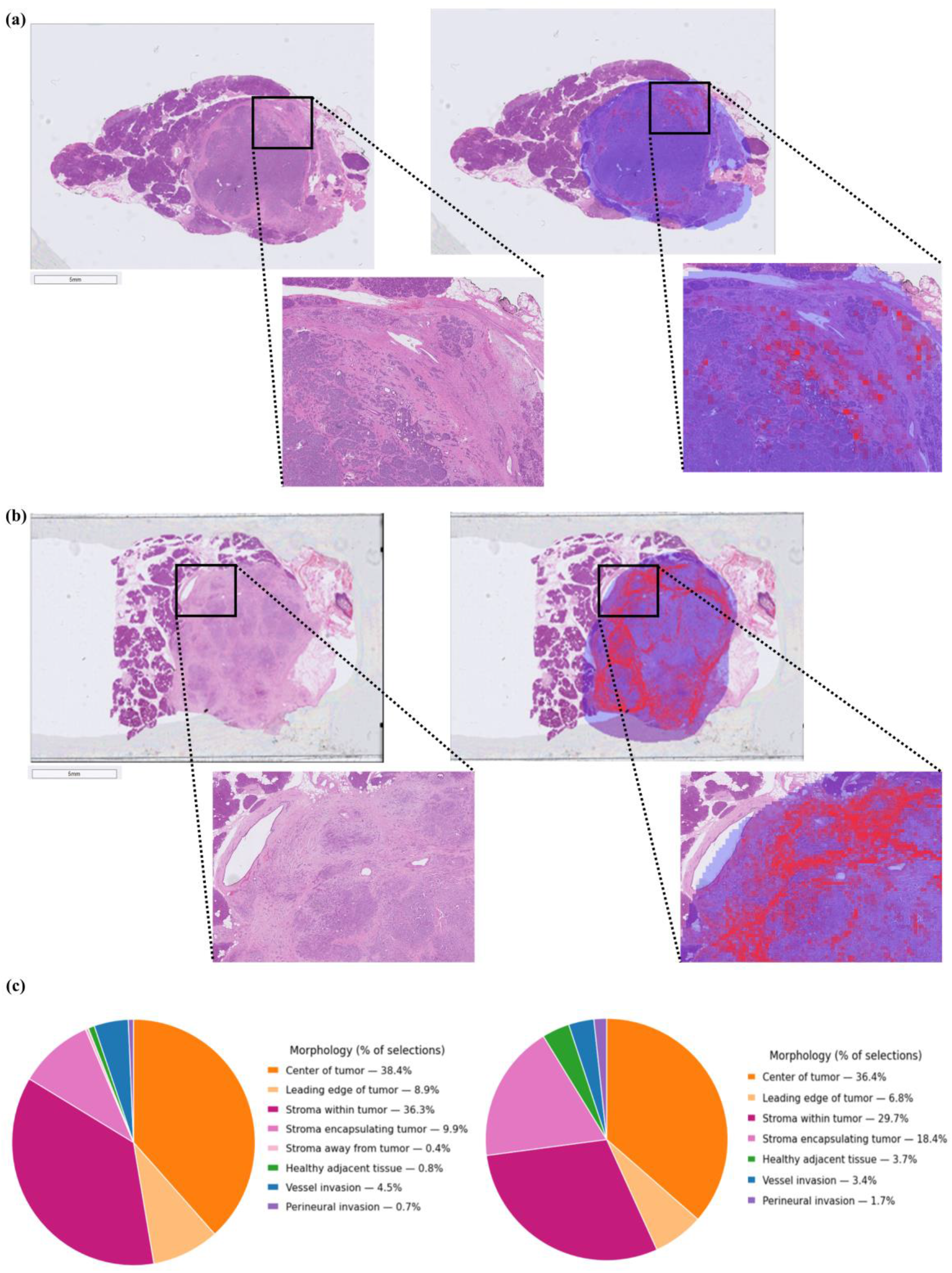
Expert review of image-model attention identifies tumor and stromal regions. **a,b**, Representative whole-slide images, attention overlays and enlarged regions showing attention at the tumor leading edge (**a**) and within intratumoral stroma (**b**). **c,** Distribution of pathologist-selected morphologic categories among reviewed regions. Intratumoral, encapsulating and tumor-distant stromal categories together account for 46.6% and 48.1% of selections in the eligible and localized cohorts, respectively.

### Multimodal integration improves or complements single-modality prediction

We next evaluated three approaches to combining structured clinicopathologic variables and primary tumor-region histology: Learned feature-level fusion, stacked Cox fusion, and fixed-weight late fusion. All used the same patient-level fold assignments as the unimodal models. Feature-level fusion jointly trained a clinical-image survival model; stacked Cox learned weights for the two unimodal risk scores; and fixed-weight late fusion averaged their standardized scores without additional outcome-based training. The late-fusion approach is illustrated in Fig. 5a, with the other approaches shown in Supplementary Fig. 4.

**Figure 5.**
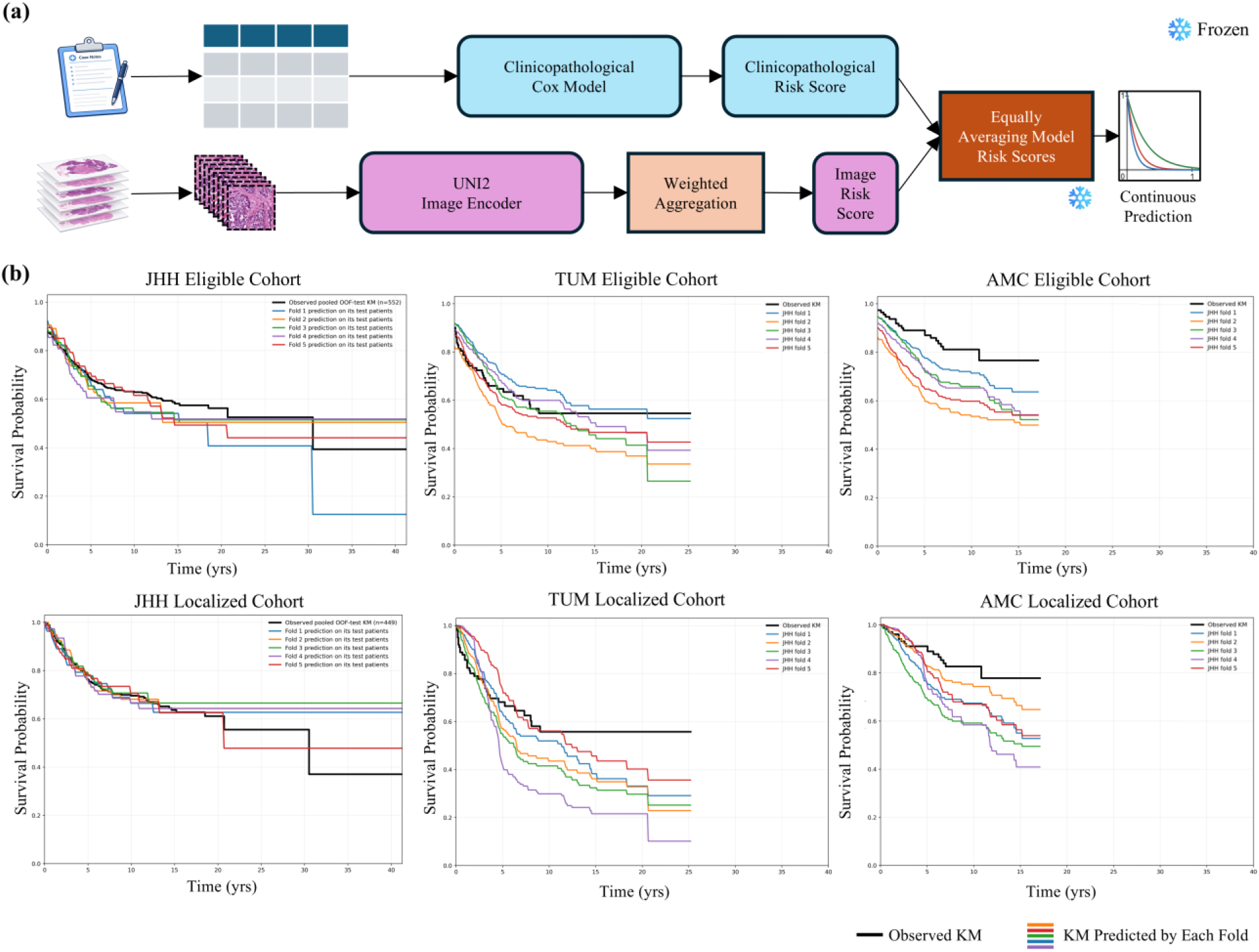
Decision-level integration of clinical and histologic survival predictions. Top, Structured clinicopathologic variables and tumor-region histology are modeled independently to generate patient-level Cox risk scores, which are combined by fixed-weight late fusion. Bottom, Observed Kaplan-Meier survival (black) and survival predicted by fivefold-specific fused models (colored) for eligible and localized cohorts at JHH, TUM and AMC. JHH predictions are from held-out test folds; each external cohort is evaluated with the five JHH-trained models.

Feature-level fusion achieved mean held-out C-indices of 0.8616 ± 0.0334 in the eligible JHH patients and 0.8385 ± 0.0196 in the localized set of patients, with corresponding five-year time-dependent AUCs of 0.9005 ± 0.0342 and 0.8737 ± 0.0378. Stacked Cox achieved C-indices of 0.8851 ± 0.0342 and 0.8360 ± 0.0309, with AUCs of 0.9124 ± 0.0345 and 0.8761 ± 0.0498, respectively. Fixed-weight late fusion achieved C-indices of 0.8838 ± 0.0356 and 0.8404 ± 0.0345, with AUCs of 0.9127 ± 0.0363 and 0.8809 ± 0.0533. Late fusion yielded the highest pooled out-of-fold C-index of the two risk-score combinations in both sets of patients: 0.8831 versus 0.8800 for stacked Cox in the eligible set of patients, and 0.8363 versus 0.8312 in the set of patients with localized disease. Its pooled five-year AUCs were 0.9142 and 0.8776, respectively. Figure 5b shows observed survival alongside survival predicted by the five-fold-specific late-fusion models.

We then applied late fusion to the AMC and TUM cohorts, including both clinical records and image embeddings, without retraining the unimodal models. For the TUM cohort, mean C-indices were 0.6817 ± 0.0604 in the eligible cohort and 0.7233 ± 0.1277 in the localized cohort, corresponding five-year AUCs were 0.6985 ± 0.1293 and 0.7575 ± 0.2466. For the AMC cohort, mean C-indices were 0.8641 ± 0.0117 in the eligible cohort and 0.8566 ± 0.0264 in the localized cohort, with AUCs of 0.8918 ± 0.0207 and 0.8500 ± 0.0088. These results suggest that the most accurate prognosticators will include a combination of the standard clinicopathologic features together with algorithms incorporating slide-level information from the patient’s primary tumor.

External means and standard deviations summarize the five JHH-trained models evaluated on the same patients at each institution. Fixed-weight fusion provided the strongest pooled internal results among the evaluated risk-score combinations, while its external discrimination varied by institution and cohort.

## DISCUSSION

This study, utilizing one of the largest multi-institutional PanNET datasets with H&E whole-slide images linked to clinicopathologic records, establishes a longitudinal multimodal framework for postoperative risk stratification in patients with surgically resected PanNETs.

Among the unimodal models, the clinical model incorporating established prognostic clinicopathologic factors achieved the highest internal discrimination, with pooled out-of-fold C-indices of 0.870 in the group of eligible patients and 0.811 in patients with a localized disease. Standardizing clinical records into 14 clinicopathologic fields allowed these factors to be evaluated consistently while retaining patients with censored follow-up. These estimates were numerically comparable to or higher than those reported in prior PanNET recurrence models, although differences in study populations and endpoints limit direct comparison.^61–64^ TNM stage was the strongest individual predictor in both the eligible and localized cohorts; tumor size and Ki-67 index also showed substantial discrimination. Removing TNM stage and Ki-67 index produced the largest mean reduction in held-out C-index in the group of all eligible patients and the patients with localized disease, respectively.

The slide-only MIL model also achieved substantial discrimination, with pooled out-of-fold C-indices of 0.798 and 0.782, demonstrating that routine H&E morphology of primary tumors contains prognostic information. These image-model results are in the range reported by histology-based survival studies, although the diseases studied and endpoints differ.^39,65–74A^ key strength of the framework is that prognostic predictions could be traced back to spatially localized regions on the original H&E slides. Tile-level attention maps enabled direct review of model-highlighted areas through a pathologist-facing interface, allowing expert assessment of whether high-attention regions corresponded to recognizable histologic compartments and adverse pathologic features. The high-attention regions in the primary tumors frequently involved tumor-stroma interfaces, leading-edge of tumor, intratumoral and peritumoral stroma, and regions compatible with vascular or perineural invasion.

Pathologists have recognized that histologic features in the tumor itself such as oncocytic, rhabdoid, clear-cell and hepatoid cytology have prognostic significance,^75–78^ however the stroma has not been equivalently emphasized as a prognosticator. Though the prominence of stromal regions in model attention is biologically plausible, given previously reported associations between fibrosis, stromal hyalinization and organization with adverse outcomes in patients with a PanNET^79–81^, the pattern is notable because contemporary PanNET risk assessment does not routinely quantify spatial organization of tumor-associated stroma and tumor-border morphology as prognosticators. Whether these patterns reflect fibrosis, fibroblast activation, extracellular-matrix remodeling or other microenvironmental changes remain to be established. These hypothesis-generating findings motivate spatial transcriptomic profiling of model-highlighted regions, paired with histologic characterization, to identify the underlying stromal programs and evaluate their potential as prognostic biomarkers.

Although image-only modeling captured substantial prognostic information, fixed-size tile embeddings may incompletely represent features such as vascular invasion that require broader tissue context. Structured clinical variables can complement these representations by retaining established prognostic findings that may be missed by the image model. Bringing the clinical and histologic predictions together yielded the strongest pooled internal discrimination: fixed-weight late fusion reached C-indices of 0.883 and 0.836 in the group of eligible patients and in the set of patients with localized disease, respectively.

External validation also exposed differences in how clinical information is recorded across institutions and over time. Variation in pathology reporting and documentation may limit the transfer of a model built from structured clinical fields. We used embedding alignment to address shifts associated with scanning and staining for the image model. These results support the feasibility of slide-based prediction across institutions, while showing that site-specific performance remains an important consideration.

This study has some limitations that should be acknowledged. The retrospective cohort spans multiple decades during which surveillance practices, pathology reporting, slide preparation, and clinical documentation evolved; scanner platforms varied across institutes; and residual heterogeneity may have influenced outcome assignment and clinical-model performance despite extensive efforts at harmonization. Furthermore, attention maps identify regions associated with model predictions but do not establish causal biology, ^82^ and cross-modal modeling was limited by the absence of direct region-level image-text supervision. Future studies should therefore validate the identified histologic phenotypes and incorporate molecular correlates, and spatial profiling to strengthen biological interpretation. Nevertheless, our findings show that long-term outcome-linked H&E histology of primary tumors contains clinically meaningful prognostic information not fully captured by conventional clinicopathologic variables and support interpretable multimodal modeling as a framework for postoperative risk stratification in patients with a surgically resected PanNETs.

## ACKNOWLEDGEMENTS

This work was supported by the Stringer Foundation and Doug and Julie Ostrover.

## CODE AVAILABILITY

Code will be made available upon peer-reviewed publication of the manuscript.

## METHODS

### Cohort design and outcome definitions

This retrospective multi-institutional study used archival surgical specimens and associated clinicopathologic and follow-up data from patients with pancreatic neuroendocrine tumors treated at JHH, USA; TUM Germany; and AMC, Republic of Korea. Patients with surgically resected pancreatic neuroendocrine tumors were identified from institutional pathology and surgical records. Available clinical case notes, pathology reports and longitudinal follow-up records were reviewed to obtain clinicopathologic characteristics, recurrence dates, vital status and cause of death where documented. Archival H&E-stained slides were retrieved and digitized as whole-slide images, with primary-tumor slides linked to each patient’s clinical data. The JHH cohort supported model development and internal cross-validation, and the TUM and AMC cohorts provided independent external validation.

### Clinical case-note standardization and text embedding

We integrated clinical and pathologic information available at surgical resection into 14 patient-level variables: age, sex, functional status (insulinoma, gastrinoma, etc.), syndromic status (multiple endocrine neoplasia 1, von Hippel-Lindau, etc.), tumor focality (unifocal/multifocal), location and size, surgical procedure, ratio of positive to harvested lymph nodes (lymph node ratio), AJCC TNM stage, Ki-67 proliferation index, vascular invasion, perineural invasion, and margin status. We standardized variable names and categorical values across records. We represented continuous variables with cubic B-spline functions to permit nonlinear associations with outcome, and one-hot encoded categorical variables. We filled missing numeric values with zero and assigned missing or out-of-definition categorical under a separate category.

We used the resulting structured features in a clinical-only Cox proportional hazards model of time to event, with censored follow-up retained in the analysis. The features were also combined with patient-level histology features in multimodal Cox models, described below. No clinical sentence templates or text embeddings were used in these survival models.

### Whole-slide image acquisition and foundation-model feature extraction

For the JHH cohort, we digitized H&E-stained surgical sections from surgically resected PanNET as whole-slide images using a Hamamatsu Nanozoomer slide scanner at 20× magnification, corresponding to a resolution of 0.46 microns per pixel. Only slides containing the primary PanNET were included for modeling, ensuring that histology-derived features were extracted from diagnostically relevant primary tumor-bearing sections. Slides in the TUM cohort were scanned on an Aperio GT450 Digital Pathology Slide Scanner, and images were converted to SVG format for analysis. Slides from the AMC cohort were scanned on a Philips IntelliSite Ultra Fast Scanner (Philips, Amsterdam, the Netherlands), and images were acquired in the proprietary iSyntax format and converted to TIFF for analysis. The external cohorts were scanned at a resolution of 0.5034 and 0.25 microns per pixel and therefore needed embedding alignment to adjust for domain shifts.

Because whole-slide images contain large areas of non-tumor pancreas, background, stroma, fat, and adjacent tissue, image modeling was restricted to tumor-centered regions of interest rather than all tiles of the entire slide. Initial tumor regions were identified using pathologist-annotated PanNET-containing areas on each WSI. To refine these annotations, we applied the CODAVision tissue segmentation framework to generate tissue-class predictions at the pixel level across the slide. The pathologist annotation and the CODAVision mask with tiles containing 10% non-tissue area were combined by taking their intersection, resulting in a high-confidence tumor RoI. The final tumor RoI was dilated by five tile widths before tile extraction to include the immediate tumor-adjacent microenvironment. This expanded RoI allowed the image model to capture not only the neoplastic cells, but also the surrounding anatomical and microenvironmental features such as invasive borders, stromal interfaces, and adjacent pancreatic tissue.

For the image model, tiles extracted from the final tumor ROI were embedded using the UNI2 image foundation model. Each image tile was encoded as a 1536-dimensional feature vector, and all tumor-region tile embeddings from a patient were grouped to form a *H_i_ ∊ R^n_i_×1536^* patient-level histology bag where *n* is the number of tiles in the RoI in patient *i*.

### Attention-based MIL

For image-only survival modeling, all UNI2 tile embeddings from the tumor region of a patient’s slides were treated as a bag of instances. Let *xij* ∈ ℝ^1536^ denote tile *j* from patient *i*, where the number of tiles *ni* varied across patients. A learned attention module assigned each tile to a score using a 128-dimensional hidden layer:

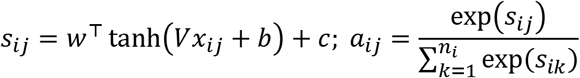

The patient-level histology representation was the attention-weighted sum *z_i_ = ∑^n_i_^_j=1_ a_ij_x_ij_*. A feed-forward network mapped *zi* to a scalar log-risk score *ri*, corresponding to the Cox proportional hazards formulation ℎ*i*(*t*) = ℎ_0_(*t*) exp(*ri*). The attention weights were retained to examine which tiles contributed most strongly to the patient-level representation.

The image model was trained on observed event times and censored follow-up using a sampled Cox partial-likelihood loss. For each patient with an observed event, the loss compared that patient’s predicted risk with the risks of up to 32 sampled patients still at risk at the event time. Separately, the structured clinicopathologic features described above were used in a clinical-only Cox model. Clinical and histology information were subsequently combined in multimodal Cox models, with the fusion procedures described in the following section.

### Multimodal survival modeling

The clinical and histology modalities were combined using three survival-modeling strategies. All approaches used the same patient-level train, validation, and test assignments as the unimodal models. Let *r^C^_i_* and *r^H^_i_* denote the log-risk scores produced by the fitted clinical Cox and histology MIL models for patient *i*, respectively.

#### a) Fixed-weight late fusion

Within each outer fold, clinical and histology risk scores were standardized separately using their means and standard deviations among that fold’s validation patients. For modality (*m* ∈ *C*, *H*), the standardized score was:

The same validation-derived transformation was applied to test-patient scores. We then assigned equal, prespecified weights to the two modalities:

This approach combined the predictions of the previously fitted unimodal models without fitting additional fusion parameters or using validation outcomes to choose the weights.

#### b) Stacked Cox fusion

A second approach fitted a Cox proportional hazards model with the two standardized unimodal risk scores as predictors:

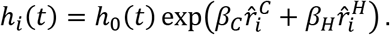

For each outer fold, the score standardization and the two Cox coefficients were estimated using that fold’s validation patients; coefficient estimation used their observed event times and censoring indicators. The stacking model used an *L*_2_ penalizer of 0.1. Its fitted coefficients were applied to the corresponding held-out test patients, while the underlying clinical and image models remained fixed. Thus, stacked Cox learned how to weight the two unimodal risk scores without retraining either modality-specific model.

#### c) Learned feature-level fusion

The third approach trained a multimodal model directly from clinical features and histology tile embeddings. Within each outer fold, the 14 clinical variables were transformed using a preprocessing pipeline fitted on the training patients, yielding a clinical feature vector *c_i_ ∊ R^d_c_^*. A trainable attention module pooled the frozen, 1,536-dimensional UNI2 tile embeddings into one histology vector *z^H^_i_ ∊ R^1536^*. The two vectors were concatenated and passed to a fusion network:

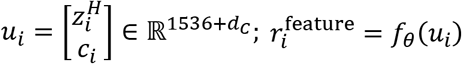

The fusion network contained hidden layers of 256 and 64 units and produced a single Cox log-risk score. Its attention module and fusion network were trained jointly using the sampled Cox partial-likelihood loss described in the training section, with up to 32 at-risk comparators per observed training event. UNI2 remained frozen. One multimodal model was trained per outer fold, and the checkpoint with the highest validation C-index was applied to that fold’s held-out test patients.

### Time-to-event modeling and Cox risk

For patient *i*, let *Ti* denote the observed time from surgery to an event or the end of follow-up, and let *δi* = 1 indicate an observed event and *δi* = 0 indicate censored follow-up. We modeled these outcomes using the Cox proportional hazards formulation. Each model assigned the patient a continuous log-risk score *ri*, which relates to the instantaneous event hazard as:

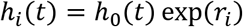

where ℎ_0_(*t*) is the baseline hazard. A higher *ri* therefore indicates a higher predicted event hazard. The source of this score differed by model: the clinical Cox model calculated it from structured clinicopathologic features, the image MIL model learned it from tumor-region tile embeddings, and the multimodal models combined clinical and histologic information.

Cox models learn relative risk from observed time-to-event while retaining censored patients in the sets of patients still at risk. For an event at *Ti*, the patient’s score is compared with the scores of patients in the risk set ℛ(*Ti*) = *j* ≥ *Ti*. The negative log partial likelihood is

The image MIL and learned feature-fusion models used the sampled-risk-set version of this objective described below. For survival-curve visualization, a fold-specific baseline cumulative hazard *H*^_0_(*t*) was estimated by the reference patients using the Breslow estimator. Predicted survival for patient *i* was then calculated as:

Patient-level risk scores were evaluated on held-out patients using Harrell’s C-index and five-year time-dependent AUC. Predicted survival curves summarized the model estimates over follow-up; observed survival was estimated separately using the Kaplan-Meier method.

### Internal cross-validation and training setup

Clinical and image survival models were evaluated using the same patient-level splits within each cohort. A five-fold stratified partition assigned each patient to one held-out test fold, approximately 20% of the cohort, with stratification based on whether an event was observed. Within the remaining patients for each fold, a fixed validation set was selected by stratified sampling of 20% of that development set. This produced approximately 64% training, 16% validation, and 20% test patients per fold, with similar observed-event proportions in the validation and test sets. The assignments were generated with a fixed random seed and shared across modalities. All slides and embeddings from a patient remained in the same partition, and each patient contributed a test prediction from one-fold.

The clinical-only model was fitted on the training patients using a Cox proportional hazards partial likelihood with an *L*_2_ penalizer of 0.1. Clinical imputation, spline transformation, scaling, and one-hot encoding were fitted on the training partition of each fold and then applied to its validation and test partitions. The penalizer was fixed across folds; the validation patients were not used to select a clinical-model hyperparameter or fit its coefficients.

The image-only model produced one log-risk score per patient from the attention-pooled tile embeddings. It was optimized using a sampled Cox partial-likelihood loss. For an observed event in patient *i* at time *ti*,up to 32 other training patients with follow-up times at least *ti* were sampled from the risk set. If *R_i_*^∗^ denotes those sampled patients, the contribution to the loss was:

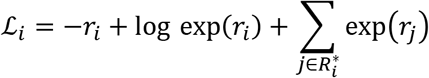

where *r* is the predicted log-risk. Patients with censored follow-up could therefore contribute as risk-set comparators while they remained under observation. The image model was trained for 30 epochs with Adam at a learning rate of 10^−3^; the checkpoint with the highest validation C-index was retained for each fold. Held-out test patients were evaluated after checkpoint selection. Performance was summarized using Harrell’s C-index and the five-year, inverse-probability-of-censoring-weighted time-dependent AUC.

Models were implemented in Python using scikit-learn and lifelines for clinical Cox modeling and PyTorch for image-model training. Image training was performed on a workstation equipped with an NVIDIA RTX 5090 GPU.

### External validation and Embedding Alignment

External validation used independent PanNET cohorts from the TUM and AMC. For clinical inference, variables in the external records were mapped to the same 14 structured fields used in the JHH cohort and processed with the saved, fold-specific clinical preprocessing pipelines. Each external patient was evaluated using the previously trained JHH models without model fine-tuning. External survival discrimination was assessed from observed event times and censored follow-up using Harrell’s C-index and the five-year time-dependent AUC.

To present tissue at a comparable physical scale to the image model (since WSIs were scanned in different scanners across institutions, and the resolutions differ), regions from each external slide were sampled with the scanner-specific pixel size accounted for, corrected for the same microns per pixel (MPP), resized to 224 × 224 pixels, and embedded with the same frozen UNI2 encoder used for the JHH slides. Tile embeddings from each patient were then grouped into a histology bag for MIL inference.

The external UNI2 embedding distributions differed from the JHH distribution (Supplementary Fig. X). Scanner characteristics, slide preparation and H&E staining, and differences in cohort composition could all contribute to this shift. We applied feature-wise location and scale alignment before image-model inference. For an external embedding component *x*ext, with external cohort’s mean and standard deviation *μ*ext and *σ*ext, and corresponding JHH reference statistics *μ*JHU and *σ*JHU, the aligned value was:

Embedding distributions before and after alignment were visualized using two-dimensional t-SNE projections (Supplementary Fig. 2).

## SUPPLEMENTARY FIGURES

**Supplementary Fig 1.**
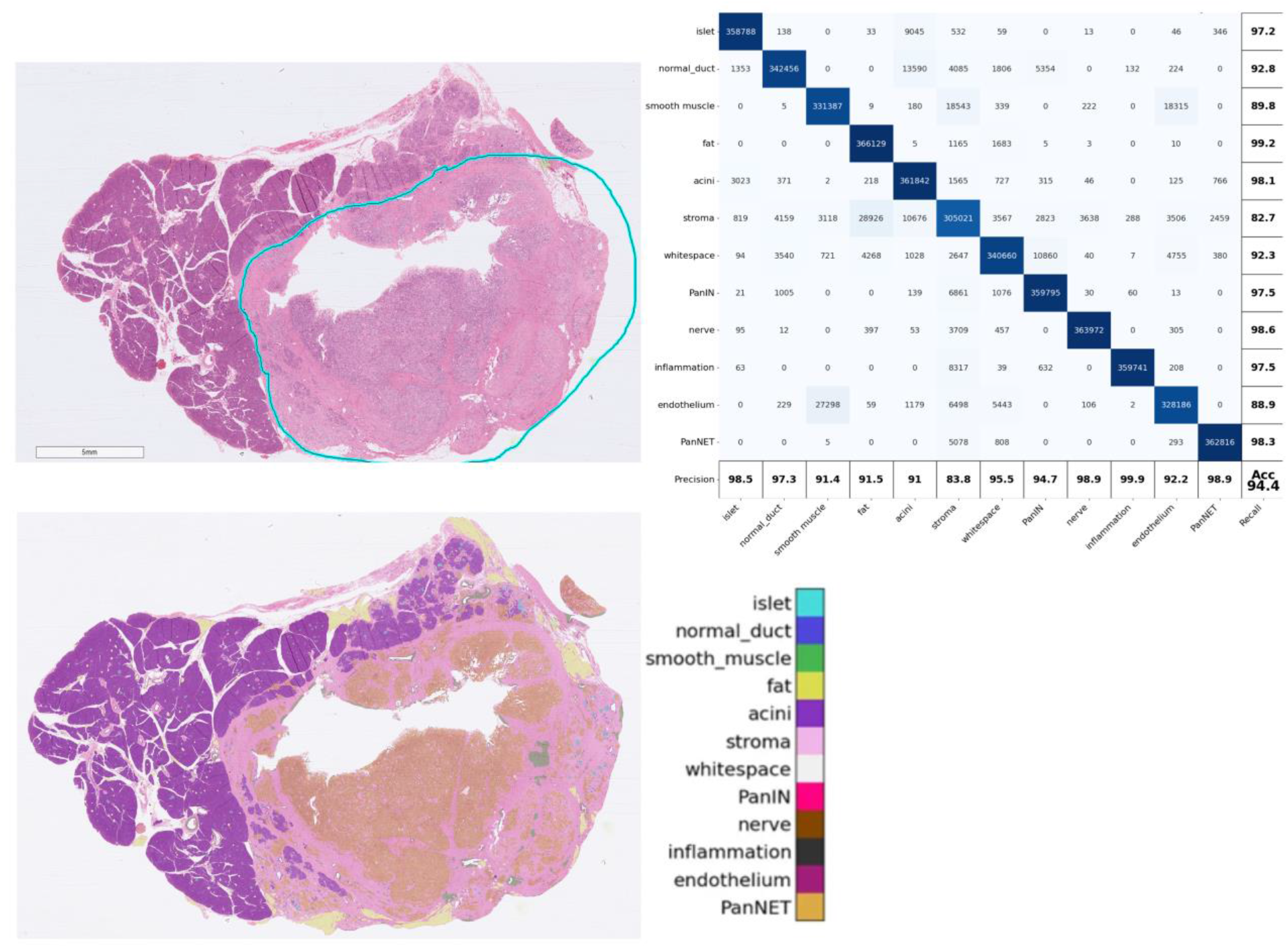
CODAVision tissue segmentation and tumor-region localization.

**Supplementary Fig 2.**
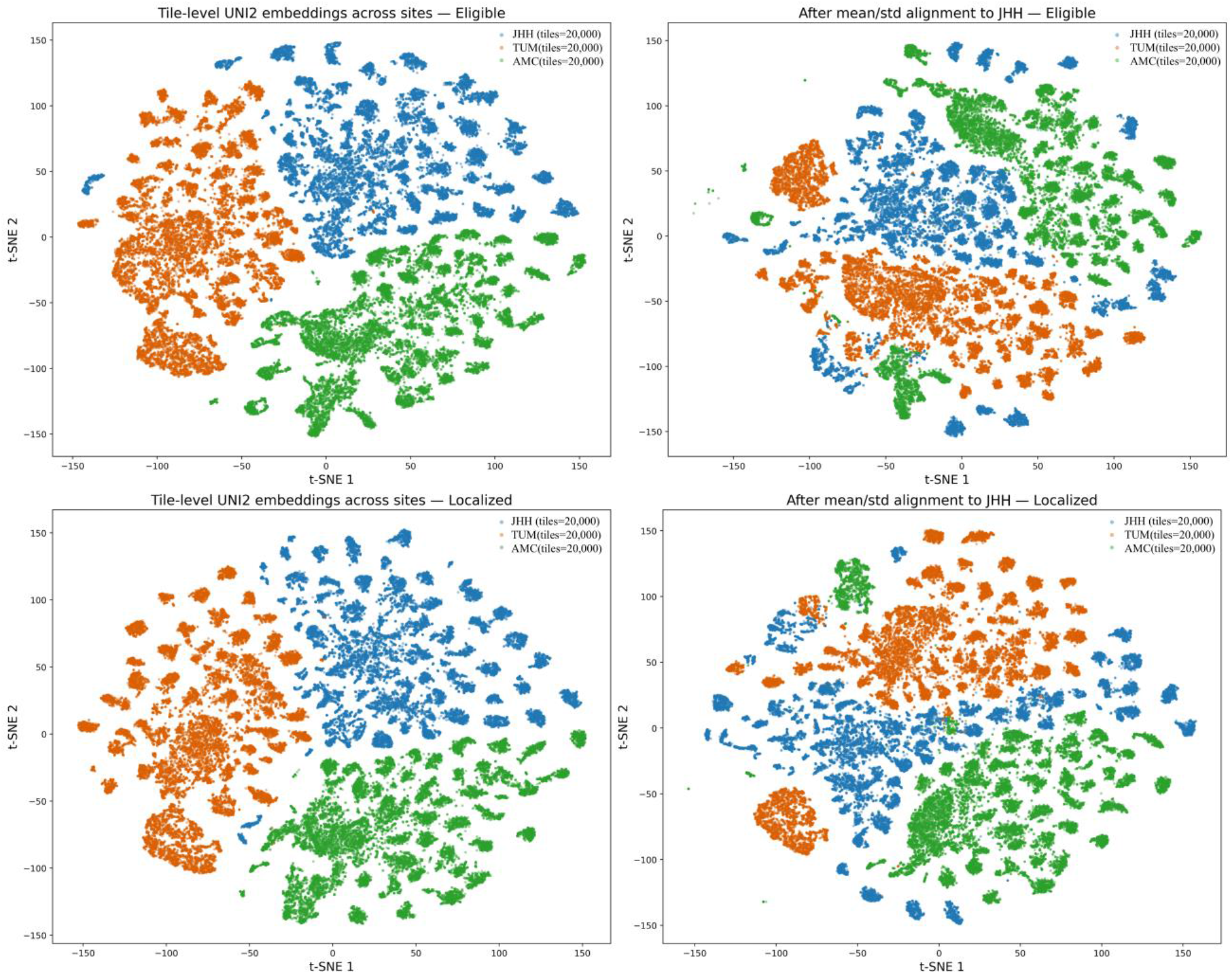
UNI2 embedding distributions across institutions before and after alignment.

**Supplementary Figure 3.**
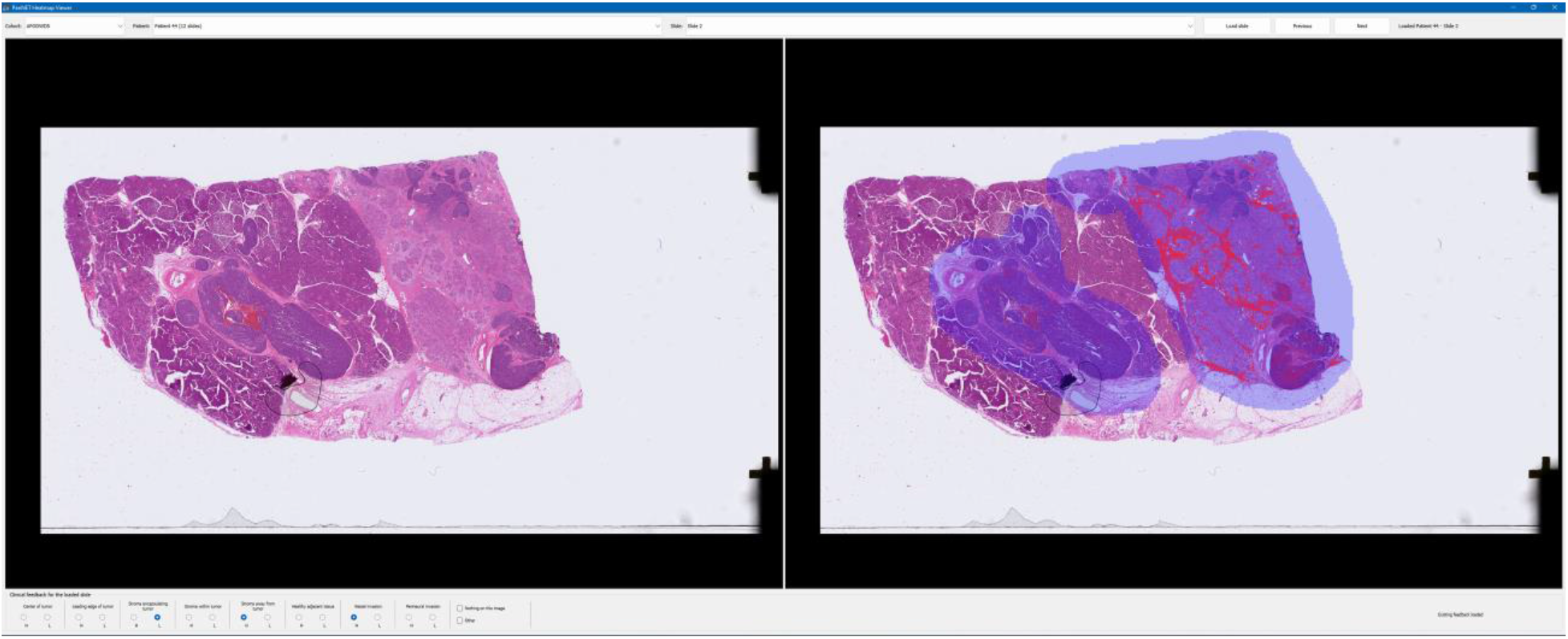
PanNET Histology and Attention Review Graphic User Interface

**Supplementary Figure 4.**
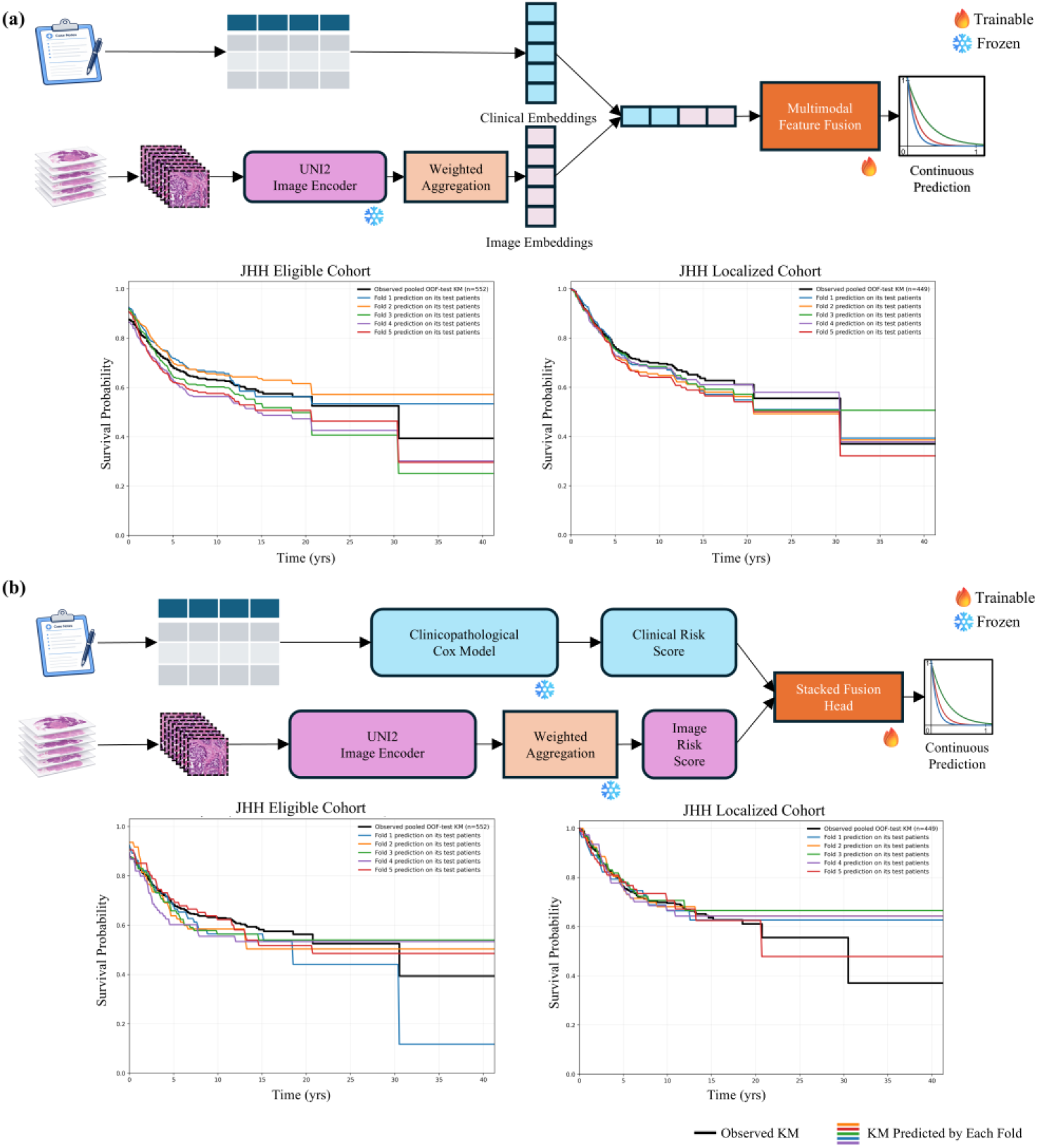
Architectures and survival curves of Multimodal Fusion and Stacked Fusion.

